# Stable reference genes for 24-hour circadian profiling of core clock genes in the blood of obstructive sleep apnea patients

**DOI:** 10.1101/2025.09.22.677736

**Authors:** Katarina Nahtigal, Ana Halužan Vasle, Tinkara Kreft, Cene Skubic, Miha Mraz, Miha Moškon, Leja Dolenc Grošelj, Damjana Rozman

## Abstract

Obstructive sleep apnea (OSA), one of the most common breathing disorders during sleep, is associated with circadian rhythm disruption, yet circadian genes expressed in blood remain largely unexplored as biomarkers. Reliable circadian gene expression analysis requires stable reference genes for qRT-PCR normalization, but no clear guidance exists for blood-based circadian studies in OSA. We evaluated the expression of 11 candidate reference genes (*GAPDH, ACTB, RPL13A, PPIB, TBP, HPRT1, PPIA, SDHA, TUBB2A, UBC*) in whole blood collected every 6 hours over 24 hours from 40 participants who underwent overnight respiratory polygraphy. Gene stability was assessed with RefFinder and EndoGeneAnalyzer across OSA severity levels, intra-individual variability, and time-point variability. *ACTB* and *RPL13A* consistently emerged as the most stable reference genes under all conditions. Their robustness was confirmed by personalized cosinor analysis of core clock genes (*BMAL1, PER2, CRY1*), showing that reliable normalization enables detection of circadian oscillations in clinical samples. While population-level analysis revealed no significant rhythmicity, individual profiles showed oscillations in 25 participants, independent of OSA severity. These findings identify *ACTB* and *RPL13A* as strong candidates for circadian transcriptomic studies in blood and provide a methodological foundation for biomarker discovery in sleep medicine, molecular medicine, and translational chronobiology.

## Introduction

Despite technological development, quantitative reverse transcription polymerase chain reaction (qRT-PCR) remains the gold standard for low throughput gene expression analysis. The accuracy of results relies heavily on the amount and quality of isolated RNA, reverse transcription efficiency, PCR amplification, and particularly of the use of appropriate reference genes, which serve as a normalization control^[1]^. The suitability of the reference (housekeeping) genes is context-dependent and the primary requirement for circadian rhythm studies is a stable 24-hour expression profile of a reference gene. This stability is difficult to achieve because the internal circadian clock influences nearly every physiological process, including the expression of reference genes themselves.

Circadian rhythm is an adaptation of living organisms to the natural cycle of environmental light and darkness, with an intrinsic period of approximately 24 hours^[2,3]^. On a molecular level, circadian rhythm is controlled by the transcriptional and translational feedback loops of core clock components. The central clock involves transcription factors (e.g., BMAL1 and CLOCK) activating clock-controlled genes (*PER* and *CRY* gene families), which in turn inhibit the activators and produce self-sustained oscillations^[2]^. This biological clock is intrinsic to almost all life forms and has gained increasing interest in the scientific community, particularly regarding its genetic foundations and impact on physiology. It is known that circadian rhythms influence different physiological processes, including metabolism, regulation of body temperature, endocrine secretion, and even labor onset^[4]^. Understanding the genetic mechanisms that drive circadian rhythms is crucial, as disruptions in these rhythms are linked to various health conditions^[5]^, including metabolic disorders (e.g., non-alcoholic fatty liver disease)^[6]^, cardiovascular diseases (e.g., heart failure)^[7]^, and sleep disturbances (e.g., obstructive sleep apnea)^[8]^.

Given the complexity and variability in gene expression driven by circadian rhythms^[9–14]^, we aim to identify reliable circadian biomarkers in the blood cells of patients with obstructive sleep apnea (OSA). OSA is a condition characterized by repeated interruptions in breathing during sleep, leading to fragmented sleep, intermittent hypoxia, and various adverse health outcomes^[2,15]^. Additionally, it has been shown that OSA has several overlaps with disrupted circadian rhythms^[3]^. It is hypothesized that the severity of OSA may correlate with distinct patterns in circadian gene expression^[2]^. A prerequisite for this investigation is the identification of stable reference genes from human blood cells for normalizing the circadian gene expression^[9,12]^. Currently, such reference genes are not well-established and predominantly based on rodents, presenting a significant barrier to accurate and reproducible research^[10,11,13,14]^. Therefore, the primary goal of this study was to determine which reference genes exhibit the most stable expression profiles across different time points and different severities of OSA. Here, we present a 24-hour analysis of candidate reference gene expression in the buffy-coat (the leukocyte-thrombocyte layer) of blood samples from patients with varying degrees of OSA. Our results identify two reference genes that are most suitable for normalizing circadian expression of core clock genes, paving the foundation for future circadian biomarker discovery.

## Materials and methods

### Subjects and blood sample collection

We included 40 participants (16 women and 24 men, age 18-65 years) with clinically suspected OSA who underwent overnight polygraphy at the Clinical Institute of Clinical Neurophysiology, University Medical Centre in Ljubljana, Slovenia. The study was approved by the Commission for Medical Ethics of the Republic of Slovenia (approval number 0120-65/2023/3), therefore all experiments were performed in accordance with its guidelines and regulations. All participants were of the same ethnic background and provided written informed consent, which was obtained in the presence of the responsible medical practitioner. Based on the apnea-hypopnea index (AHI) obtained from overnight polygraphy (Alice NightOne, Philips), participants were classified into four groups: control (AHI <5), mild (AHI 5-14.9), moderate (AHI 15-29.9), and severe (AHI ≥30) OSA. Each group consisted of 10 individuals.

The peripheral venous blood was collected through peripheral venous catheter in 3 mL tubes containing EDTA anticoagulant every 6 hours throughout a 24-hour period (T0 at 13:00, T1 at 19:00, T2 at 1:00, T3 at 7:00, and T4 at 13:00 o’clock the next day, denoted as T0, T1, T2, T3, T4, respectively). After every blood withdraw within a 30-minute time window, 300 μL thrombocyte-leukocyte layer (buffy coat) was extracted from whole blood after the centrifugation of samples (15 min, 2000 × g, 4 °C) and removal of the plasma layer. The buffy coat samples were processed for downstream RNA isolation.

### RNA isolation, cDNA preparation and qRT-PCR analysis

Total RNA was extracted using TRI Reagent LS (Sigma-Aldrich). The RNA isolation protocol is described in the Supplements. RNA quantity and purity were assessed by spectrophotometry (NanoDrop 1000, Thermo Scientific). We used QuantiTect Reverse Transcription kit (Qiagen) for reverse transcription. A qRT-PCR analysis was then carried out using PowerTrack SYBR Green Master Mix (Applied Biosystems) on a QuantStudio 5 instrument (Applied Biosystems), following the standard operating procedure of manufacturer’s cycling conditions. All reactions were performed in triplicates and included no-template controls to check for contamination. PCR amplification specificity was confirmed by melt-curve analysis, ensuring a single peak for each primer set.

### Primer design

For the candidate reference genes, we selected 11 most common housekeeping (reference) genes: *GAPDH, SDHA, PPIB, ACTB, CDK4, HPRT1, RPL13A, TUBB2A, PPIA, UBC, TBP*. This selection was based on literature research^[1,7,9,14,16–21]^ and previous studies^[10]^ of our research group that highlighted these genes as potential housekeeping controls. Where available, we included multiple transcript variants or distinct primer sets for certain genes (denoted as “#2”) to assess alternative splice variants or amplicon targets (for instance, *SDHA #2, PPIB #2, CDK4 #2*). We self-designed primers using NCBI Primer-BLAST, adhering to stringent criteria to ensure efficient and specific amplification. The primer design requirements are summarized in Supplementary Table S1. We tested amplification efficiency and specificity of all primer pairs. Primer efficiency for each target was evaluated by standard curve analysis (dilution series) in qRT-PCR, and only primers with efficiency between ∼90-110% were used. Specificity was further confirmed by gel electrophoresis of PCR products, which yielded single bands of expected size. Primer sequences, amplicon lengths, and amplification efficiencies are listed in Supplementary Table S2. All qRT-PCR data (quantification cycle values) were collected and managed using QuantStudio design and analysis software version 1.6.1.

### Data analysis

Raw quantification cycle (Cq) values from qRT-PCR were used to assess reference gene expression levels. We employed multiple algorithmic approaches to determine the optimal reference genes and the minimum number of references needed for reliable normalization. We analysed raw Cq values with two different algorithm softwares used for comprehensive stability analysis and compared the results. RefFinder^[22]^ is an online tool for the evaluation of housekeeping genes, which integrates four established computational programs – geNorm^[23]^, NormFinder^[24]^, BestKeeper^[25]^, and comparative Delta Ct (ΔCt) method^[26]^. It ranks candidate reference genes from most stable to least stable. RefFinder calculates a geometric mean of acquired stabilities for each gene. This provides an overall final ranking and gives a straightforward result of best performing reference gene (lower score indicates higher stability)^[22]^. The online tool EndoGeneAnalyzer, on the other hand, allows for more complex experimental designs: it can handle multiple grouping factors (e.g., different patient groups, time points), performs inter-group comparisons, removes outliers, and computes stability metrics (based on NormFinder) within each subgroup or condition^[27]^. This enabled us to evaluate reference gene performance not only globally but also within specific conditions such as different OSA severity levels and intra-individual time-course variation.

For reference gene selection, we followed a stepwise elimination approach guided by the analysis outcomes. Finally, to validate the usefulness of the chosen reference genes, we analyzed the rhythmic expression of key clock genes (*BMAL1, PER2*, and *CRY1*) in our sample set. Cosine curve fitting was performed using CosinorPy, a Python-based implementation of cosinor rhythmometry, to detect and characterize 24-hour oscillations in gene expression^[28]^. For the cosinor analysis, we normalized *BMAL1, PER2*, and *CRY1* Cq values using the best-performing reference genes identified in this study.

## Results

### Reference gene selection process

40 participants (16 women and 24 men) aged 18 to 65 years, with suspected OSA were enrolled for reference gene analysis. They were allocated into control, mild, moderate, and severe OSA severity groups based on overnight polygraphy results of AHI. Each group consisted of 10 participants. We divided the study into three major steps, which are graphically shown in Fig. 1, to progressively narrow down the candidate reference genes.

**Figure 1:**
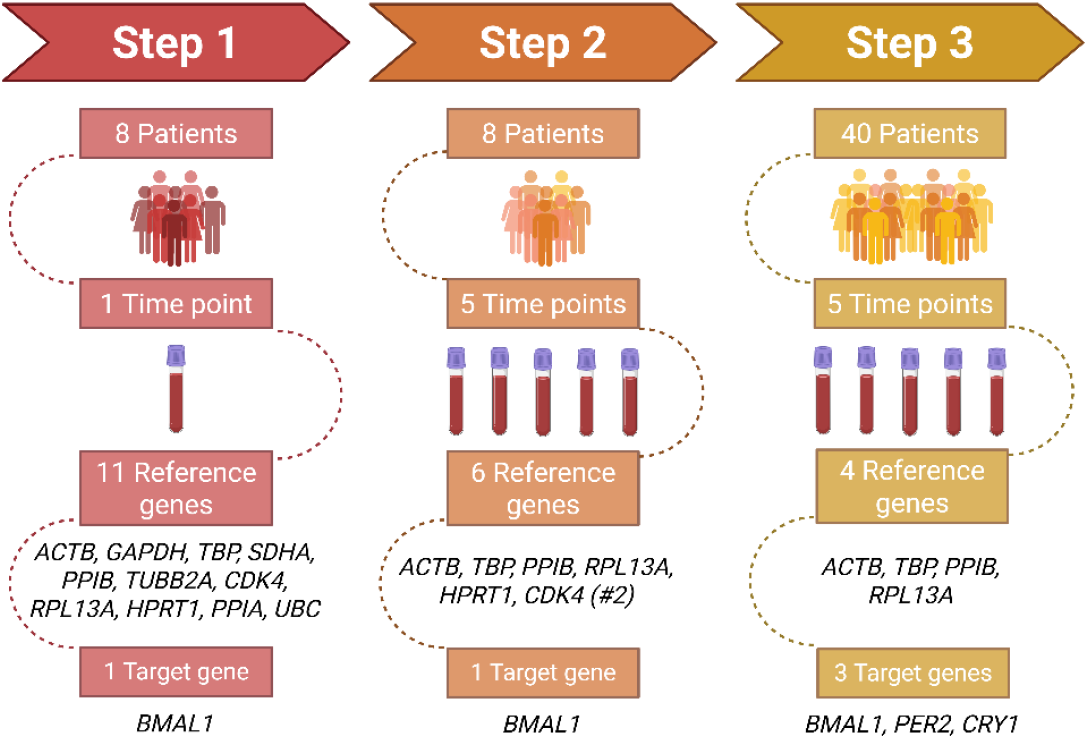
Schematic overview of the three-step reference gene selection process in this study. The study was conducted in three sequential steps. In each step, we increased the scope of analysis (either the number of participants, the number of blood sampling time points, and/or the number of target genes) while correspondingly narrowing down the list of candidate reference genes. At each step, the elimination of reference gene candidates was based on the stability results obtained with algorithms implemented in online tools RefFinder and EndoGeneAnalyzer. Step 1 involved 8 participants sampled at a single time-point to evaluate the expression stability of 11 candidate reference genes, using *BMAL1* as a representative target gene. In step 2, we used the same 8 patients and included samples from all 5 time-points to assess circadian expression patterns and refine the selection to the 6 most stable reference genes. In step 3, the final analysis was expanded on a cohort of 40 patients, sampled at 5 time-points, using the 4 most stable reference genes (*ACTB, TBP, PPIB, RPL13A*) from previous steps for normalisation. Expression of three core clock genes (*BMAL2, PER2*, and *CRY1*) was evaluated in this final phase. “#2” denotes an alternative primer sequence targeting a different transcript of the same gene.

#### Step 1

We examined the expression stability of 11 candidate reference genes (*GAPDH, SDHA, PPIB, ACTB, CDK4, HPRT1, RPL13A, TUBB2A, PPIA, UBC, TBP*), three additional alternative sequences (*SDHA #2, PPIB #2, CDK4 #2*), and one core clock gene (*BMAL1*) in a plot set of 8 participants (2 per each OSA severity group) at a single time point (at 13:00, corresponding to T0). In this initial screening, since samples were all taken at the same circadian time, we grouped data only by OSA severity condition. We analysed Cq values with both RefFinder and EndoGeneAnalyzer. Because we considered only one condition, we based our selection process more on the results obtained from RefFinder, since it is more established. Based on the stability rankings, we eliminated 6 genes (*UBC, TUBB2A, GAPDH, SDHA*, and *PPIA*; plus the *SDHA #2* and *PPIB #2* variants) that showed the highest variability (i.e., high standard deviation or poor stability scores) in this step. The detailed stability values from each algorithm are provided in Supplementary Figs. S1–S2 and Supplementary Tables S3–S4.

#### Step 2

We carried forward the 6 best-performing reference gene candidates from Step 1 (*PPIB, ACTB, CDK4 #2, HPRT1, RPL13A, TBP*), and evaluated their expression in the same 8 participants across all time points (T0-T4 over 24 h). This allowed us to assess within-individual circadian stability as well as between-group differences. We included *BMAL1* as a target gene in EndoGeneAnalyzer to help evaluate how reference gene choice might affect detection of a known rhythmic gene. In Step 2, we analysed the data in three ways: (a) grouping by individual (to assess variation within each person’s 24 h profile), (b) grouping by OSA severity (to assess differences among the four patient groups), and (c) grouping by time point (to assess overall circadian fluctuation). EndoGeneAnalyzer was particularly useful here, as it calculated stability metrics within each grouping condition. The results (Supplementary Fig. S3 and Supplementary Table S5) showed that some genes had consistently low stability in all scenarios. In particular, *HPRT1* and *CDK4 #2* emerged as less stable (higher variability) across conditions. RefFinder (Supplementary Fig. S4 and Supplementary Table S6) also placed these two genes in mid-to-low stability range. Therefore, we eliminated *HPRT1* and *CDK4 #2* at this stage.

#### Step 3

In the final step, we expanded the analysis to 40 participants and focused on the top four reference genes from step 2 (*PPIB, ACTB, RPL13A, TBP*). We measured their expression at all 5 time points in every participant. Additionally, we included three circadian target genes (*BMAL1, CRY1, PER2*) to enable downstream rhythmicity analysis. We repeated the multi-condition stability analysis with EndoGeneAnalyzer, examining each of the three factors (individual patient variation, OSA severity, and time of day) separately. Figure 2 summarizes the EndoGeneAnalyzer results for this step: graph (a) shows the standard deviation of each reference gene under each condition, while graph (b) shows the corresponding stability values (lower values indicate more stable expression). The more detailed results of the EndoGeneAnalyzer are presented in Supplementary Table S7.

**Figure 2:**
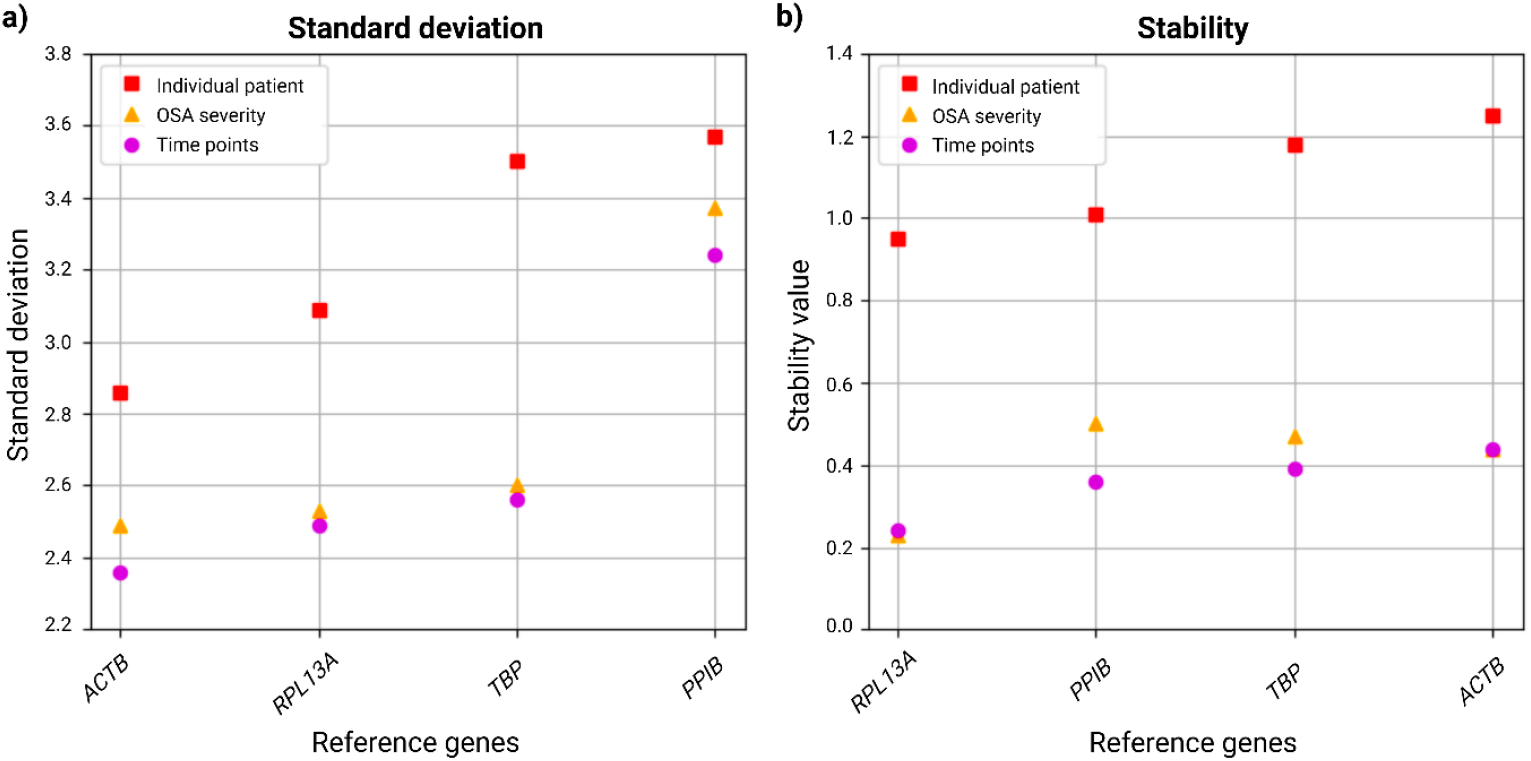
EndoGeneAnalyzer results of the four candidate reference genes in Step 3. Standard deviation (a) and stability (b) of the four best-performing reference genes (*ACTB, RPL13A, TBP, PPIB*) are shown based on three grouping criteria: individual patient (red squares), OSA severity (orange triangles), and time points (purple circles). Lower standard deviation and stability values indicate more consistent gene expression. *RPL13A* consistently exhibited the lowest variability (a) and highest stability (b) across all grouping conditions, followed by *ACTB* and *TBP*.

The EndoGeneAnalyzer analysis of Step 3 demonstrated *ACTB* having the lowest standard deviation levels throughout all three conditions, followed by *RPL13A* being second best (Fig. 2a). In addition, *RPL13A* also showed the lowest stability level in all three conditions, however, *ACTB* did not have the best stability (Fig. 2b).

Similarly, we analysed the Step 3 data with RefFinder, and the aggregated stability outcomes are presented in Supplementary Table S8. The RefFinder results are visualised in Fig. 3, which integrates the rankings from BestKeeper (Fig. 3a), geNorm (Fig. 3b), NormFinder (Fig. 3c), Delta Ct method (Fig. 3d), and the overall comprehensive score (Fig. 3e).

**Figure 3:**
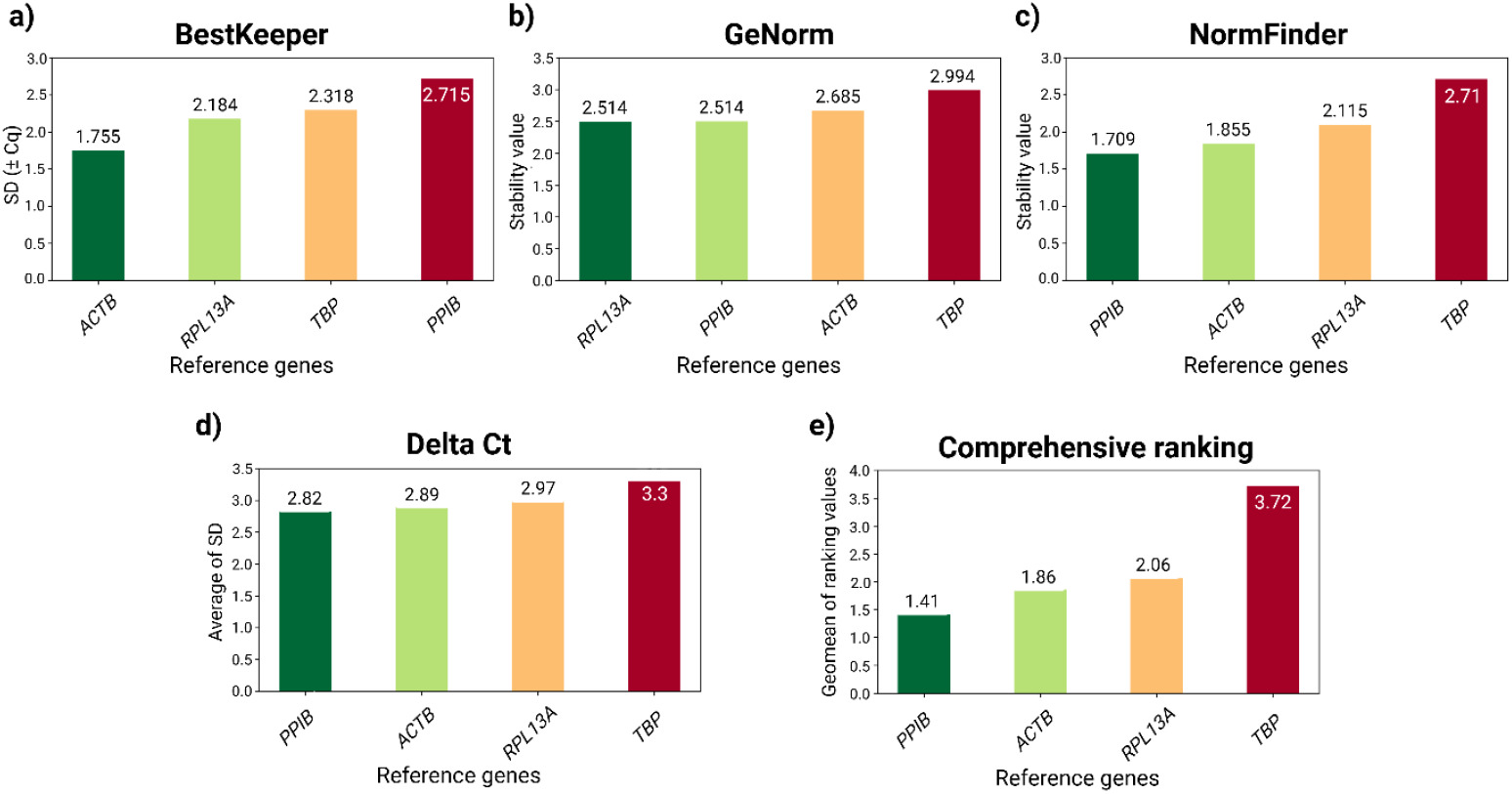
RefFinder analysis of reference gene stability in Step 3. The figure shows results from four algorithms (BestKeeper, geNorm, NormFinder, and the Delta Ct method) with additional comprehensive ranking. Each graph (a-e) represents one analysis method, plotting the stability or variability measures for *ACTB, RPL13A, PPIB*, and *TBP*. In all cases, lower values (left side) correspond to greater expression stability. *ACTB* was identified as the top performer by BestKeeper (a), *RPL13A* ranked highest by geNorm (b), and *PPIB* showed the best stability by NormFinder (c), lowest average of SD by ΔCt method (d) and overall best geometric mean of ranking values by comprehensive ranking (e). *TBP* consistently showed the lowest stability by all algorithms (a-e).

In the RefFinder analysis of Step 3, *ACTB* and *RPL13A* consistently ranked among more stable genes. For instance, *ACTB* was identified as top reference by BestKeeper (Fig. 3a), and second best by NormFinder (Fig. 3c), Delta Ct method (Fig. 3d), and comprehensive ranking (Fig. 3e). Additionally, *PPIB* also demonstrated very low stability values by NormFinder (Fig. 3c), Delta Ct method (Fig. 3d), and comprehensive ranking (Fig. 3e). On the contrary, *TBP* demonstrated as the least stable across all algorithms used by RefFinder (Fig. 3).

After Step 3, considering the collective evidence from both software tools and all conditions tested, we selected *ACTB* and *RPL13A* as the best-performing reference genes for subsequent analyses. These two genes showed low intra- and inter-group variability and remained relatively unchanged across the circadian cycle. Notably, *PPIB* also demonstrated favorable stability metrics in some analyses (especially in RefFinder’s comprehensive ranking). However, we encountered practical issues with *PPIB* during qRT-PCR: its expression was low, and in some samples, *PPIB* Cq values were undetectable even after repeated measurements. In contrast, *ACTB* and *RPL13A* were robustly detected in all samples on the first pass. Due to these technical considerations and *PPIB*’s slightly higher variability under certain conditions (Fig. 2a), we decided not to include *PPIB* as a reference for final normalization. *TBP* was also excluded based on its poor stability performance (it ranked lowest in all five algorithms, indicating significant expression variability; Figs. 2 and 3). In summary, *ACTB* and *RPL13A* were chosen as the optimal reference genes for circadian gene expression studies in OSA blood samples.

### Assessment of core clock genes rhythmicity

The Cq values describing the expression of core clock genes *BMAL1, CRY1*, and *PER2* were calculated using the comparative CT method, where *ACTB* and *RPL13A* were used as reference genes for normalisation^[29]^. These data were then used to reconstruct the group-specific cosinor models for each group of participants and for each gene separately. In this case, a cosinor model is built for each individual and each gene. The models describing the same gene and individuals from the same group are averaged to obtain a group-specific model (Fig. 4). The obtained models were then used to assess the rhythmicity parameters of each group (Supplementary Table S9).

**Figure 4:**
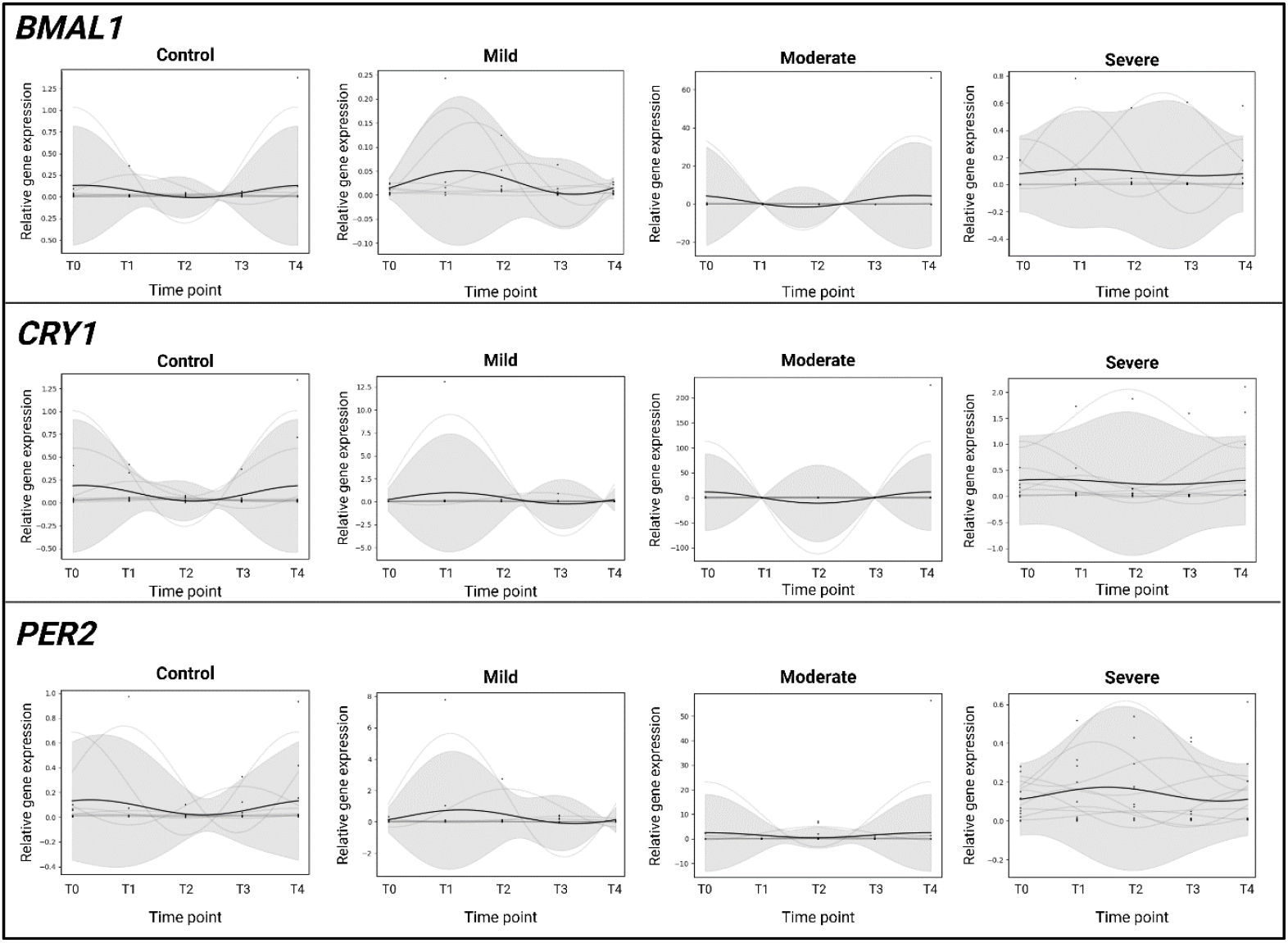
The circadian expression profiles of core clock genes (*BMAL1, CRY1*, and *PER2*) across OSA severity groups. Gene expression rhythms were assessed using a population-based cosinor for control, mild, moderate, and severe OSA groups. Cq values were normalized using *ACTB* and *RPL13A* as reference genes. Each subplot represents the fitted group-specific rhythm (black line), obtained by averaging individual-specific cosinor models (grey lines). Shaded areas denote 95% confidence intervals. No statistically significant rhythmicity was observed across severity groups for any of the genes analysed. Sampling time (x-axis): T0 on day 1 at 13:00, T1 on day 1 at 19:00, T2 on day 2 at 1:00, T3 on day 2 at 7:00, and T4 on day 2 at 13:00.

Group-specific cosinor analysis revealed that none of the core clock genes exhibited statistically significant 24-hour rhythmicity across the OSA severity groups. For *BMAL1*, rhythmicity was absent, and amplitudes remained low (0.024–3.08). Interestingly, both mild OSA (20.84) and severe OSA (20.5) showed later acrophases. The mesor values for *BMAL1* were also modest, ranging from 0.026 to 1.43, without consistent differences across groups. Similarly, *CRY1* demonstrated low to moderate amplitudes across groups (0.05– 11.20), with the largest amplitude observed in the moderate OSA group (11.20). Statistical tests for amplitude again did not achieve significance in any group. Mesor values ranged from 0.10 to 0.36, where mild OSA showed slightly higher values (0.358) compared to controls (0.102). Acrophase values varied between 13.93 h and 19.99 h, across groups, though none reached statistical significance and rhythmicity remained non-significant. The gene *PER2* exhibited low amplitudes (0.036–1.02) across all groups, indicating no significant rhythmicity. Amplitudes were highest in the moderate OSA group (11.02) but failed to reach statistical significance. Mesor values ranged from 0.08 to 1.48, where moderate OSA group showed the highest value (1.48), however with no significant group differences (Supplementary Table S9).

Although the results of cosinor analysis of the investigated core clock genes did not show statistically significant 24-hour rhythmicity of the observed genes when 10 patients in the group were considered, the rhythmicity of each individual patient and individual-specific models are still apparent in Fig. 4. Closer look into the oscillation patterns of each individual patient reveals severe desynchronization within different OSA severity group and clock genes presented in Fig. 5.

**Figure 5:**
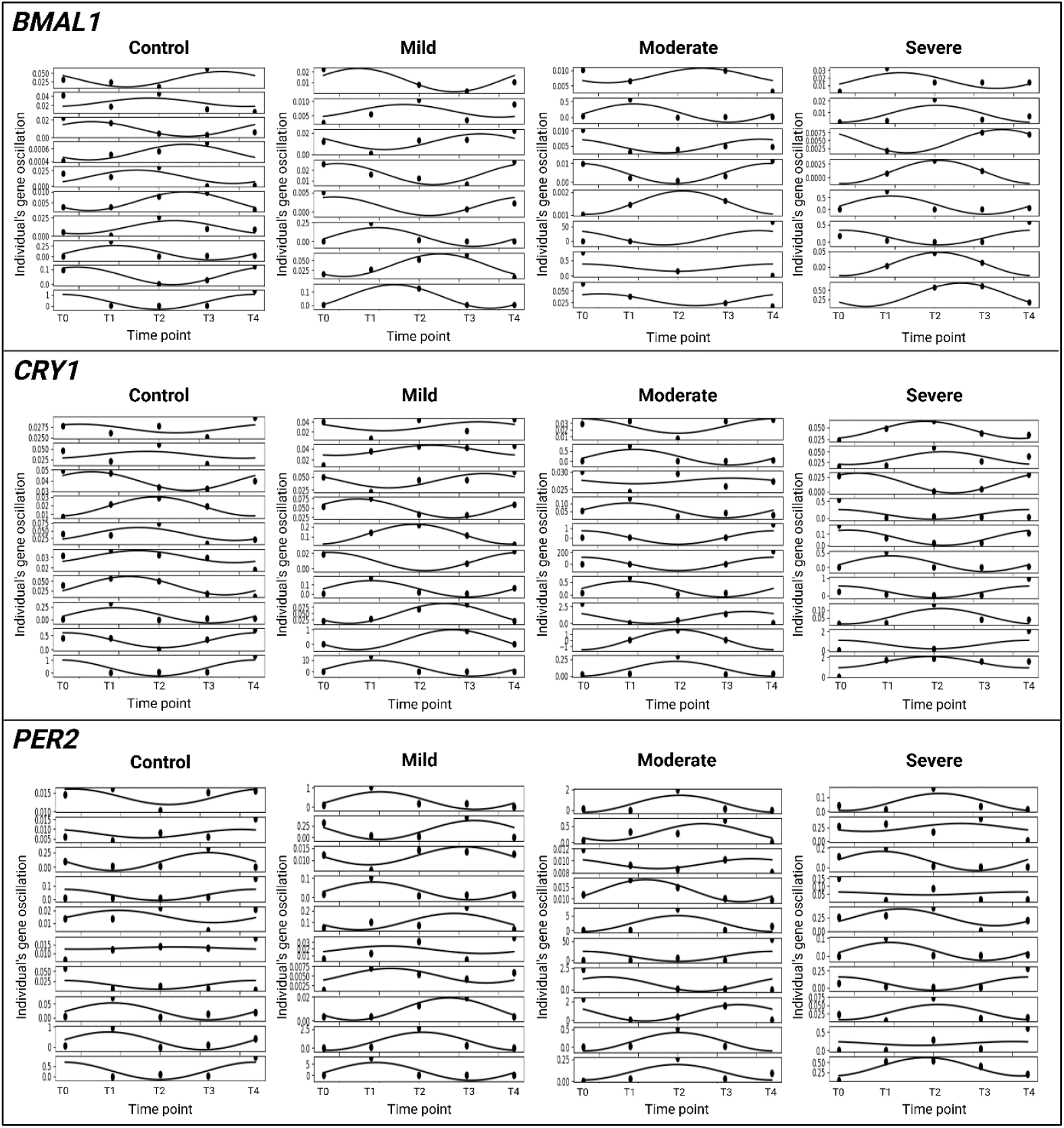
The circadian expression of core clock genes (*BMAL1, CRY1*, and *PER2*) of individual participants within each OSA severity group obtained from population-based cosinor analysis. Each panel represents the fitted cosinor curves (black lines) and raw expression data (black dots) for each individual participant. Rows correspond to different target genes, while columns correspond to OSA severity groups. Timepoints (x-axis) are plotted over a 24-hour period, and the y-axis represents normalised gene expression levels. Population-based cosinor was used to estimate individual mesor, amplitude, and acrophase parameters, highlighting inter-individual variability in circadian rhythmicity within each group. This approach reveals both well-defined oscillations and flattened or phase-shifted profiles, underscoring the heterogeneity of circadian rhythms in patients with OSA. Sampling time (x-axis): T0 on day 1 at 13:00, T1 on day 1 at 19:00, T2 on day 2 at 1:00, T3 on day 2 at 7:00, and T4 on day 2 at 13:00.

Because population-based models did not show significant 24-hour rhythmicity of the observed genes in groups of 10 participants, we also performed the personalized cosinor analysis, which shows clear oscillatory patterns in several patients, emphasizing inter-individual variability, which may be independent of the OSA severity. This underlines the importance of patient-specific circadian assessment (Fig. 6; Supplementary Table S10).

**Figure 6:**
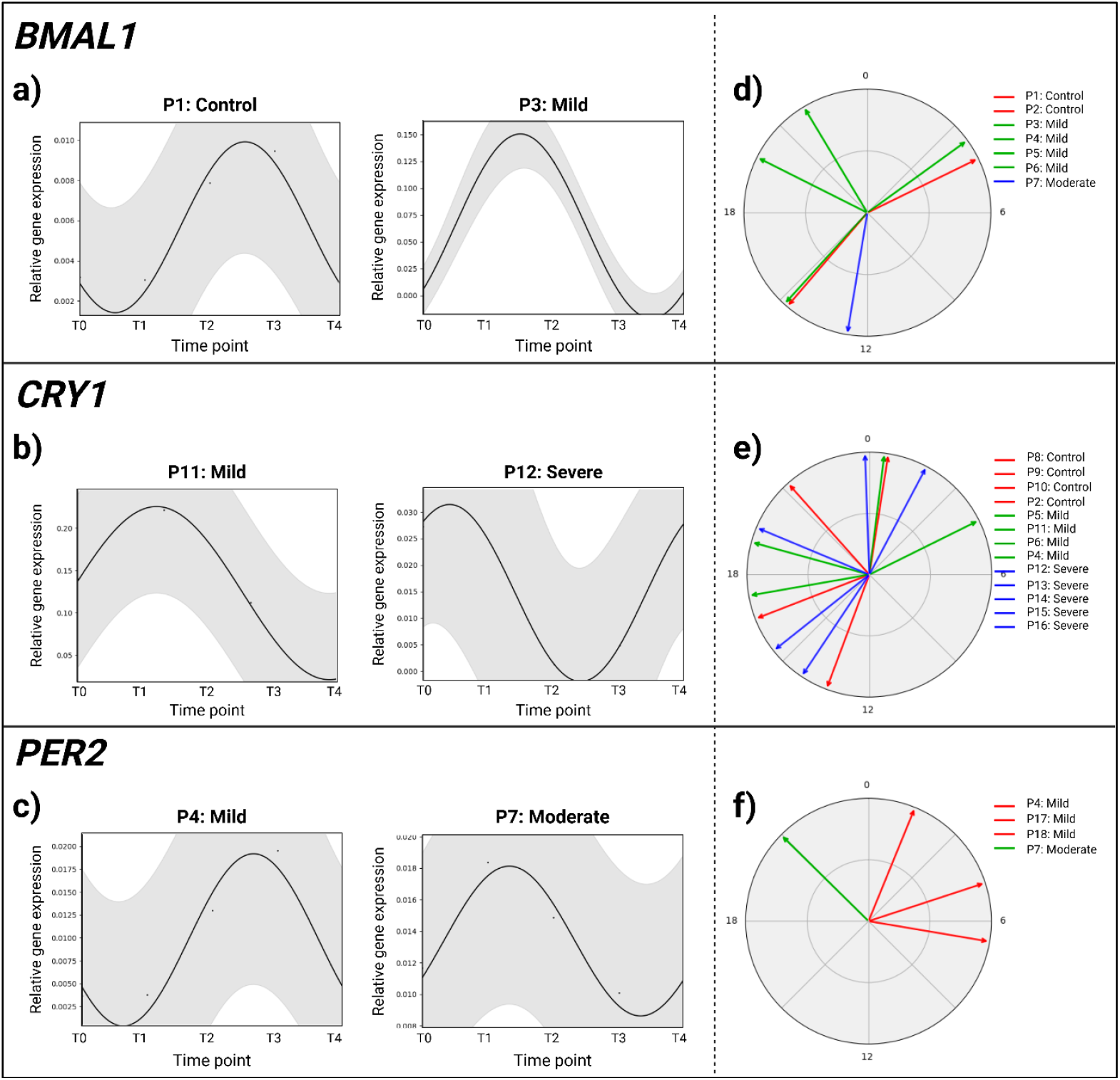
Personalized cosinor analysis of core clock genes in OSA patients and controls. Panels a–c show individual cosinor fits (black lines) with 95% confidence intervals (grey areas) for representative participants, illustrating gene-specific oscillatory patterns of *BMAL1* (a), *CRY1* (b), and *PER2* (c). Each subplot corresponds to one participant, where “P” followed by a number denotes a specific individual (e.g., P1 = Participant 1). Panels d–f display acrophase distributions derived from CosinorPy for *BMAL1* (d), *CRY1* (e), and *PER2* (f). Each line represents different participant denoted with P and a number. Sampling time (x-axis): T0 on day 1 at 13:00, T1 on day 1 at 19:00, T2 on day 2 at 1:00, T3 on day 2 at 7:00, and T4 on day 2 at 13:00.

The personalized analysis revealed statistically significant rhythmicity of *BMAL1* (n=7), *CRY1* (n=13), and *PER2* (n=5) in 25 participants with different OSA severity levels (Supplementary Table S11). The phase plots (Fig. 6d-f) show substantial inter-individual variability in circadian timing, with no consistent phase alignment within severity groups. Collectively, these findings highlight the heterogeneity of circadian gene expression in OSA and underscore the importance of a personalized approach in circadian rhythm assessment.

## Discussion

In this study, we addressed a critical methodological gap for circadian biology research – the selection of suitable reference genes for normalizing gene expression over the 24-hour cycle. Our approach followed a rigorous three-step selection framework, progressively narrowing down from an initial pool of 11 candidate reference genes (and 3 additional primer sequences of *CDK4, SDHA*, and *PPIB*) to the two most stable genes. Using both RefFinder and EndoGeneAnalyzer tools, we systematically identified and validated *ACTB* and *RPL13A* as the most stable reference genes in the buffy coat of human blood, under conditions that included intrapatient variability, different OSA severities, and multiple circadian time points. To our knowledge, this is the first comprehensive, time-resolved assessment of reference gene stability for circadian studies in peripheral blood from patients with OSA, enabling reliable detection of rhythmic molecular signals in clinical cohorts.

In the first screening step, candidate reference genes were assessed at a single time point using RefFinder and EndoGeneAnalyzer, which allowed efficient exclusion of the most variable transcripts before moving to more resource-intensive analyses. Extending the evaluation across five time points in Step 2, revealed marked circadian instability for *HPRT1* and *CDK4 #2*, leading to their exclusion. The final analysis in Step 3, based on 40 participants with time-series blood sampling, integrated multiple stability algorithms and consistently identified *ACTB* and *RPL13A* as the most stable reference genes across all OSA severity groups. Importantly, both genes displayed minimal diurnal variability in buffy coat, despite the well-known context-dependence of housekeeping gene expression. This context-dependence refers to the fact that housekeeping genes are not universally stable, since their expression can vary with tissue type, physiological state, or even time of day, and therefore must be validated within each experimental setting^[1,30]^.

Our findings align with previous work in clinical and circadian research where *ACTB*^[16,31–33]^ and *RPL13A*^[34,35]^ have been validated as stable internal controls in blood-derived samples^[36–38]^. However, they contrast with studies in other tissues reporting pronounced circadian oscillations of *ACTB*, such as in the mouse lung^[1]^, or under altered physiological conditions such as sleep deprivation^[30]^. These discrepancies underscore that no single gene is universally suitable as a reference and that reference gene selection must be tailored to the biological context, tissue, and experimental design. The strong performance of *RPL13A* in our analysis is consistent with prior reports suggesting that ribosomal proteins are often stably expressed under hypoxic conditions^[34,35]^. Likewise, Ledderose et al. (2011) identified *RPL13A* as a reliable reference gene in human leukocytes exposed to various stimulations, supporting its robustness across diverse immune and stress-related contexts^[11]^. By systematically validating internal controls for circadian qPCR in OSA patient blood, our study provides a methodological framework for future clinical chronobiology research.

On the contrary, several candidate genes commonly used as references were found to be unsuitable in our circadian OSA context. *GAPDH*, for example, is a classic housekeeping gene also used in circadian studies^[39]^ but was eliminated in Step 1 of our study due to high variability. This is not entirely surprising, as *GAPDH* expression can be influenced by factors like metabolic state and time of day^[40–42]^. *TBP* was another gene that performed poorly, ranking as the least stable by all algorithms in our final analysis of Step 3. Our data contradicts other studies that reported *TBP* to be a stable reference gene for circadian studies using human mammary epithelial cells^[12]^. These observations reinforce that even genes involved in basic cellular maintenance (housekeeping) are not inherently constant under all conditions and must be validated. *PPIB* was another gene that showed promising performance in the study focusing on circadian rhythms in different mouse tissues and strains^[12]^. However, despite a good stability ranking in RefFinder, *PPIB* had inconsistent amplification in some blood samples, making it a less practical choice. Such real-world factors are sometimes overlooked if one focuses purely on algorithm output, so we recommend that researchers always combine computational ranking with empirical lab observations when selecting reference genes. Additionally, recent approaches in literature recommend using multiple reference genes for normalization to improve accuracy and compensate for any minor fluctuations in one gene’s expression^[23]^. In line with those recommendations, our use of two reference genes (*ACTB* and *RPL13A*) should provide a more reliable normalization factor for circadian gene studies in OSA than any single gene alone.

In addition, an important strength of our study was the integration of different computational tools like RefFinder and EndoGeneAnalyzer to rigorously evaluate gene stability. RefFinder allowed us to integrate several established ranking methods, providing a consensus view of stability. EndoGeneAnalyzer, a recently introduced tool (2024), was particularly valuable for our complex experimental design. It enabled us to simultaneously consider multiple grouping factors (time, OSA severity, intra-individual differences) and identify reference genes that are consistently stable across all these dimensions. The concordance between the two tools in pinpointing *ACTB* and *RPL13A* increases our confidence in these genes. We also highlight that practical considerations (such as primer efficiency and gene expression level) influenced our final choice.

Using *ACTB* and *RPL13A* for normalization, we then assessed the rhythmic expression of core clock genes *BMAL1, CRY1*, and *PER2*. Surprisingly, none of these genes demonstrated statistically significant 24-hour rhythmicity in any of the OSA severity groups when using population-based cosinor model. This result diverges from findings in healthy populations, where peripheral clock gene oscillations are often observed, albeit with modest amplitudes^[16,18,31–33,43,44]^. Several factors could explain this discrepancy: (1) peripheral blood may be a less robust tissue for detecting circadian rhythms in OSA populations; (2) OSA itself may dampen or desynchronize peripheral clock gene expression due to intermittent hypoxia, sleep fragmentation, and metabolic disturbances; (3) inter-individual variability and modest sample sizes may have limited statistical power to detect subtle rhythmicity; (4) CosinorPy method requires a minimum of five time points, therefore we might lack sufficient temporal resolution to robustly detect subtle circadian patterns, especially if gene expression exhibits low amplitude or interindividual variability. Whereas, majority of studies researching circadian oscillation of clock genes is not using the same mathematical methods^[16,18,31–33,43,44]^. Though cosinor models visually suggested oscillatory trends, particularly in *CRY*1 and *PER2*, the group-level models failed to reach significance.

Nonetheless, there were subtle, non-significant trends in phase timing across severity groups. For instance, *BMAL1* exhibited slightly increased mesor in moderate OSA (1.43) and earlier acrophases (from 13.91 h in controls to 11.28 h in moderate OSA group), suggesting a possible phase advance, though not statistically significant. On the contrary, *PER2* acrophase was most delayed in the severe group (22.19 h), suggesting a trend toward phase delay in advanced OSA, albeit without statistical support (Supplementary Table S9). These trends, although not statistically significant, could point to a progressive disruption of the molecular circadian system with increasing disease severity. Further studies using higher-powered datasets and additional peripheral tissues may be required to confirm this observation.

Our personalized cosinor analysis revealed that circadian gene expression in OSA patients is highly heterogeneous, with significant oscillatory patterns detected in some individuals but not consistently across severity groups (Supplementary Tables S10 and S11). The acrophase plots (Fig. 6d–f) clearly illustrate the wide inter-individual variability, with participants displaying distinct phase distributions even within the same OSA category. Notably, *CRY1* showed the highest number of participants with significant rhythmicity (n=13), whereas *PER2* exhibited the lowest (n=5), suggesting that different clock components may be differentially affected by OSA (Supplementary Table S11). Importantly, these rhythms did not cluster according to disease severity, indicating that OSA-related circadian disruption is not solely a function of apnea severity but may be influenced by additional factors such as metabolic status, lifestyle, or genetic background^[45]^. This variability highlights the limitations of population-averaged models, which failed to detect significant rhythms, and underscores the value of personalized circadian assessment in uncovering individual-specific alterations. Together, these findings support the hypothesis that OSA exerts heterogeneous effects on circadian regulation, reinforcing the need for individualized approaches in both research and clinical chronotherapy^[46–49]^.

Our findings have significant implications for circadian rhythm research in OSA. OSA patients experience chronic intermittent hypoxia and sleep fragmentation, which are known to perturb molecular clock function. Indeed, emerging evidence indicates that OSA is associated with altered expression of core clock genes and clock-controlled genes in various tissues^[20,36,37,49]^. For example, a recent study by Turkiewicz et al. (2025) demonstrated differential expression of multiple core clock genes (*BMAL1, CLOCK, PER1, CRY1, NPAS2, NR1D1*) in OSA patients’ blood, alongside changes in inflammatory and neurotrophic signaling pathways^[50]^. Understanding these alterations could provide insight into OSA’s comorbidities (cardiovascular, metabolic, mood disorders, etc.) and identify potential biomarkers or therapeutic targets.

A key strength of this study is the integration of multiple reference gene validation tools and the consideration of circadian and disease-related grouping variables in stability analysis. This represents a rigorous approach to normalization in time-series gene expression studies. However, some limitations must be acknowledged. The lack of rhythm in core clock genes could partly result from the use of a specific blood cell fraction (buffy coat), which contains a heterogeneous mix of cell types. While blood is a practical tissue for chronobiology studies and buffy coat contains relevant immune cells that can reflect circadian patterns, it is possible that different tissues or different patient demographics would have other optimal reference genes. Future studies may benefit from sorting specific immune cell populations or using RNA sequencing to assess clock gene expression in greater depth. However, our aim was to use the most robust and easily accessible material for investigating gene expression variability throughout the 24-hour circadian rhythm.

Additionally, the sample size (40 subjects) for gene expression analysis may limit statistical power, especially in detecting modest rhythmic patterns. Larger cohorts and longitudinal designs, especially pre- and post-treatment, could clarify whether OSA interventions (e.g., CPAP therapy) restore molecular rhythmicity and provide even more power to detect subtle expression variability.

Overall, our multi-step approach and consistency across methods give us results that emphasize the importance of rigorous reference gene validation when studying circadian transcription in clinical populations. The identification of *ACTB* and *RPL13A* as stable reference genes for diurnal qPCR studies in patients with OSA provides a valuable foundation for future research. We also note that using two reference genes in combination (as we recommend) involves more experimental effort than a single reference, but the improvement in data reliability is well worth it despite the increase in cost and labor. Moreover, the observed discordance between molecular and hormonal circadian outputs underscores the complexity of circadian dysregulation in OSA. Further studies with larger cohorts, transcriptomic-scale data, and interventional designs (e.g., CPAP treatment) will be necessary to clarify whether peripheral clock gene dampening is a feature or consequence of OSA, and how it relates to clinical outcomes.

## Acknowledgments

The authors used ChatGPT (OpenAI) for AI assisted copy editing for text improvements, readability, grammar corrections, and spelling. All content generated with its help was carefully reviewed and edited by all the authors, who take full responsibility for the final published version. The figures in the manuscript were edited in the BioRender.com online tool. This research study is funded by Slovenian Research Agency (ARIS) P1-0390, P2-0359, P3-0338, and J1-50024 grants, the infrastructure ESFRI-ELIXIR grant, and a tertiary project of UMC Ljubljana No. 20220001. The authors would like to thank medical student Lucija Matijašić and Hana Rakuša, M.D., for their valuable work, sleepless nights, and assistance in this study.

## Author contributions statement

K.N. study management and administration, conceptualization, validation, investigation, conceived and conducted the experiments, data analysis, and drafted the manuscript. A.H.V. and M. Moškon data analysis, methodology. T.K. and C.S. conceptualization, methodology. L.D.G. was the responsible physician of the clinical study. M. Mraz, L.D.G. and D.R. conceptualization and supervision. All authors edited and reviewed the final manuscript.

## Data availability statement

Data supporting this study are included in the manuscript and supplementary files. Additional datasets are available from the corresponding author upon reasonable request, in accordance with privacy and ethical regulations.

