## Supplements for "Stable reference genes for 24-hour circadian profiling of core clock genes in the blood of obstructive sleep apnea patients"

### RNA isolation protocol

Total RNA was extracted using TRI Reagent LS (Sigma-Aldrich) in a 1:1 ratio to the volume of thrombocyte-leukocyte layer samples. The samples were stored at -80 °C overnight. The next day, phase separation was done following the manufacturer’s protocol. RNA participation was done by adding ice-cold isopropanol in a 1:1 ratio to the volume of the transferred aqueous phase. After washing the samples with 75% ethanol solution, we stored the samples at -80 °C overnight. The following day, we washed RNA two more times with a 75% ethanol solution. The RNA solubilisation was performed by drying the samples for approximately 20 minutes at room temperature and adding of appropriate volume (app. 20 µL) of RNA-free water to the RNA pellet. Then the samples were incubated for 5 minutes at 55 °C.
