## Supplementary Fig. S1 for "Stable reference genes for 24-hour circadian profiling of core clock genes in the blood of obstructive sleep apnea patients"

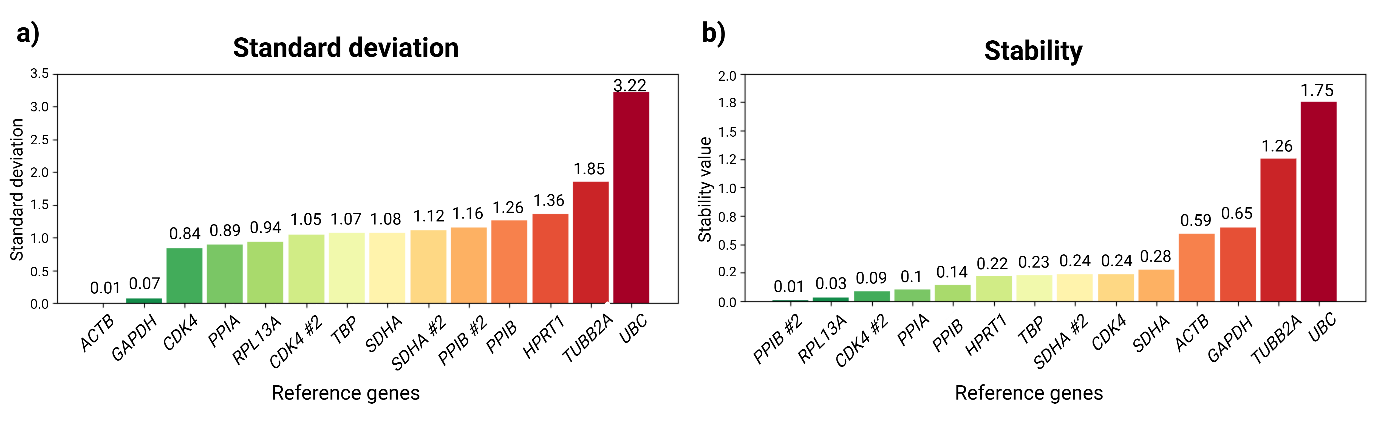

**Supplementary Figure S1:** **Results of step 1 candidate reference genes using EndoGeneAnalyzer**. Reference gene stability was assessed based on (a) standard deviation and (b) stability value across all samples. Lower values indicate higher expression stability. Genes are ranked from most (left side) to least (right side) stable. The “#2” label denotes an alternative primer sequence.
