## Supplementary Fig. S2 for "Stable reference genes for 24-hour circadian profiling of core clock genes in the blood of obstructive sleep apnea patients"

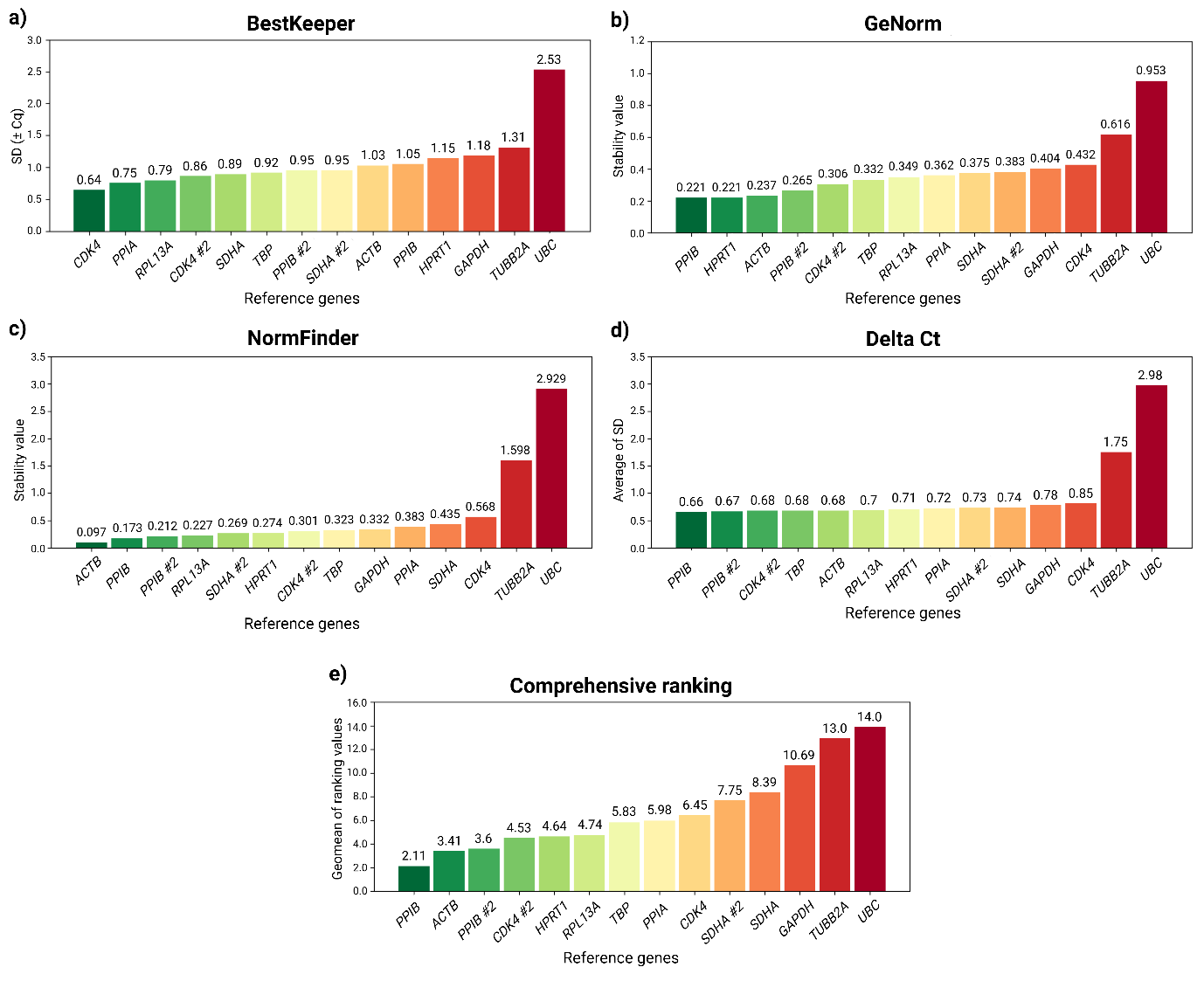

**Supplementary Figure S2:** **Stability evaluation of candidate reference genes using RefFinder based on Step 1 data**. RefFinder integrates multiple statistical algorithms to assess the stability of reference genes. Panels show (a) standard deviation (SD ± Cq) values calculated by BestKeeper, (b) stability values from GeNorm, (c) NormFinder, and (d) the Delta Ct method. Lower values indicate greater gene expression stability across samples. (e) A comprehensive ranking summarizes results from all four algorithms, where lower stability values indicate higher overall gene stability. The “#2” label indicates an alternative primer sequence.
