## Supplementary Fig. S3 for "Stable reference genes for 24-hour circadian profiling of core clock genes in the blood of obstructive sleep apnea patients"

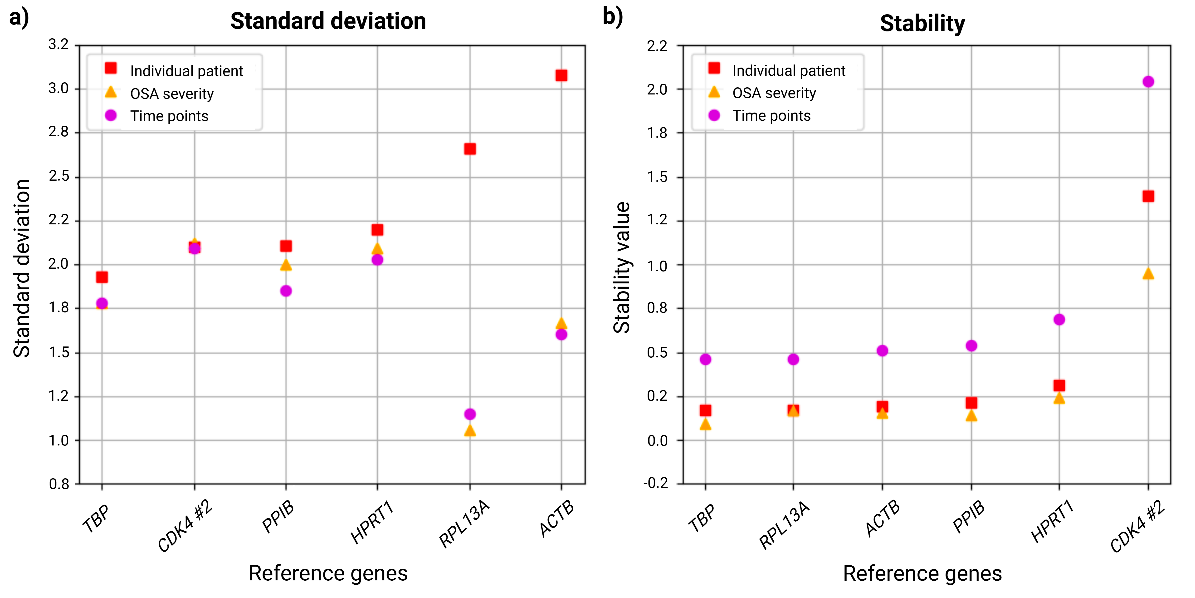

**Supplementary Figure S3:** **Evaluation of step 2 reference gene performance under different conditions using EndoGeneAnalyzer.** (a) Standard deviations and (b) stability values of candidate reference genes were assessed across three conditions: individual patients (red square), OSA severity groups (orange triangles), and time points (purple circles). Lower values indicate greater expression stability. “#2” indicates an alternative primer sequence.
