## Supplementary Fig. S4 for "Stable reference genes for 24-hour circadian profiling of core clock genes in the blood of obstructive sleep apnea patients"

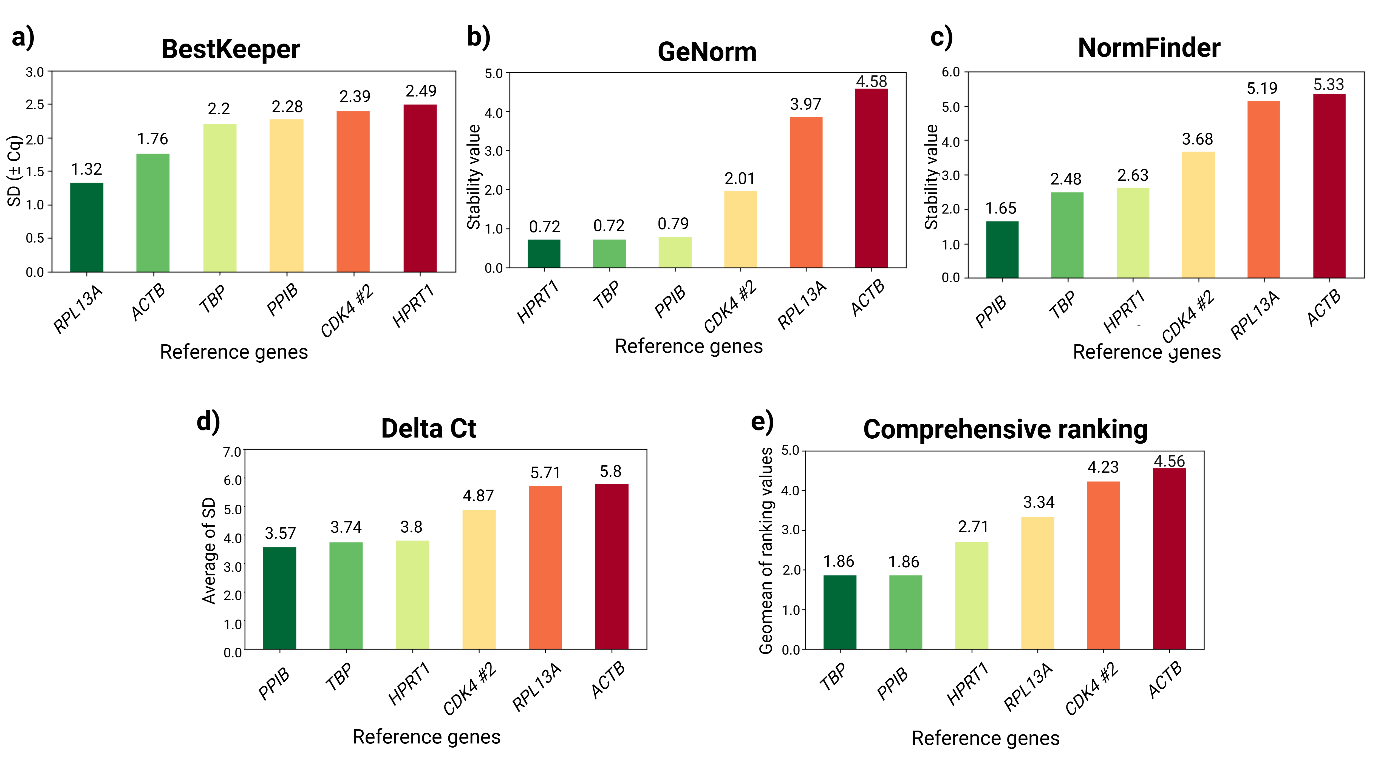

**Supplementary Figure S4:** **Comparative assessment of reference gene stability using RefFinder based on data from Step 2.** The stability of candidate reference genes was evaluated using four different algorithms: (a) standard deviation calculated by BestKeeper, (b) stability ranking by GeNorm, (c) NormFinder, (d) Delta Ct method, and (e) comprehensive ranking which integrates results from all four methods. Lower values (left side) indicate greater gene expression stability. “#2” indicates an alternative primer sequence.
