## Supplementary Table S1 for "Stable reference genes for 24-hour circadian profiling of core clock genes in the blood of obstructive sleep apnea patients"

**Supplementary Table S1:** qRT-PCR primer design criteria.

| **PCR product size:** | from 70 to 200 bp |
| --- | --- |
| **Number of primers to return:** | 10 |
| **Tm:** | min = 57 °C, max = 63 °C, opt. = 60 °C |
| **Exon junction span:** | must span an exon-exon junction |
| **Organism:** | 9606 (human) |
| **Tm difference between forward and reverse sequence:** | max 3 °C |
| **GC percentage:** | 40-60% |
| The lowest value of self-complementary. | |
| The lowest number of potentially unintended templates. | |
