## Supplementary Table S2 for "Stable reference genes for 24-hour circadian profiling of core clock genes in the blood of obstructive sleep apnea patients"

**Supplementary Table S2:** Primer sequences and efficiency of reference genes.

| **Symbol** | **Gene name** | **NCBI Gene ID** | **Primer sequence (5’ – 3’)** | **Amplicon length (bp)** | **Amplification efficiency (%)** |
| --- | --- | --- | --- | --- | --- |
| ***ACTB*** | Actin, beta | 60 | FW: ACAGAGCCTCGCCTTTGCC  RV: GATATCATCATCCATGGTGAGCTGG | 70 | 101 |
| ***BMAL1*** | Basic helix-loop-helix ARNT like 1 | 406 | FW: GCTCAGGAGAACCCAGGTTATC  RV: GCATCTGCTTCCAAGAGGCTCA | 161 | 100 |
| ***CDK4*** | Cyclin dependent kinase 4 | 1019 | FW: GTGTATGGGGCCGTAGGAAC  RV: GATCAAGGGAGACCCTCACG | 89 | 96 |
| ***CDK4 #2*** | Cyclin dependent kinase 4, second primer sequence | 1019 | FW: CCATCAGCACAGTTCGTGAGGT  RV: TCAGTTCGGGATGTGGCACAGA | 103 | 95 |
| ***CRY1*** | Cryptochrome circadian regulator 1 | 1407 | FW: GCAGTTGCTTGCTTCCTGACAC  RV: GACAGCCACATCCAACTTCCAG | 125 | 93 |
| ***GAPDH*** | Glyceraldehyde-3-phosphate dehydrogenase | 2597 | FW: TGGAAGGACTCATGACCACA  RV: TTCCCGTTCAGCTCAGGGAT | 169 | 100 |
| ***HPRT1*** | Hypoxanthine phosphoribosyl-transferase 1 | 3251 | FW: TGCTTTCCTTGGTCAGGCAG  RV: TTCAAATCCAACAAAGTCTGGC | 110 | 106 |
| ***PER2*** | Period circadian regulator 2 | 8864 | FW: AGCTGCTTGGACAGCGTCATCA  RV: CCTTCCGCTTATCACTGGACCT | 118 | 85 |
| ***PPIA*** | Peptidylprolyl isomerase A | 5478 | FW: GCCAAGACTGAGTGGTTGGAT  RV: GGCCTCCACAATATTCATGCC | 75 | 99 |
| ***PPIB*** | Peptidylprolyl isomerase B | 5479 | FW: TCTCCGAACGCAACATGAAG  RV: AATTCGTAGGTCAAAATACACCTTG | 146 | 97 |
| ***PPIB #2*** | Peptidylprolyl isomerase B, second primer sequence | 5479 | FW: AACGCAGGCAAAGACACCAACG  RV: TCTGTCTTGGTGCTCTCCACCT | 140 | 92 |
| ***RPL13A*** | Ribosomal protein L13a | 23521 | FW: CGCCCTACGACAAGAAAAAGC  RV: TACTTCCAGCCAACCTCGTG | 118 | 98 |
| ***SDHA*** | Succinate dehydrogenase complex flavoprotein subunit A | 6389 | FW: GTGCATTTGGTGGACAGAGC  RV: TCATATCGCAGAGACCTTCCA | 124 | 104 |
| ***SDHA #2*** | Succinate dehydrogenase complex flavoprotein subunit A, second primer sequence | 6389 | FW: GAGATGTGGTGTCTCGGTCCAT  RV: GCTGTCTCTGAAATGCCAGGCA | 142 | 95 |
| ***TBP*** | TATA box-binding protein | 6908 | FW: TGTATCCACAGTGAATCTTGGTTG  RV: GGTTCGTGGCTCTCTTATCCTC | 124 | 93 |
| ***TUBB2A*** | Tubulin beta 2A class IIa | 7280 | FW: TTGGGAGGTCATCAGCGATGAG  RV: AGGCTCCAGATCCACCAGGATG | 151 | 98 |
| ***UBC*** | Ubiquitin C | 7316 | FW: CCGGGATTTGGGTCGCAG  RV: TCACGAAGATCTGCATTGTCAAG | 70 | 100 |

* Abbreviations: NCBI – National Center for Biotechnology Information; FW – forward sequence; RV – reverse sequence.
