## Supplementary Table S3 for "Stable reference genes for 24-hour circadian profiling of core clock genes in the blood of obstructive sleep apnea patients"

**Supplementary Table S3:** Results of standard deviation and stability from EndoGeneAnalyzer, comparing different OSA severity levels from step 1 of reference gene selection.

| **Gene(s)** | **Standard deviation** | **Stability value** |
| --- | --- | --- |
| ***ACTB*** | 0.01 | 0.59 |
| ***GAPDH*** | 0.07 | 0.65 |
| ***TBP*** | 1.07 | 0.23 |
| ***TUBB2A*** | 1.85 | 1.26 |
| ***SDHA*** | 1.08 | 0.28 |
| ***SDHA* #2** | 1.12 | 0.24 |
| ***PPIB*** | 1.26 | 0.14 |
| ***PPIB* #2** | 1.16 | 0.01 |
| ***CDK4*** | 0.84 | 0.24 |
| ***CDK4* #2** | 1.05 | 0.09 |
| ***RPL13A*** | 0.94 | 0.03 |
| ***HPRT1*** | 1.36 | 0.22 |
| ***PPIA*** | 0.89 | 0.10 |
| ***UBC*** | 3.22 | 1.75 |

*Abbreviation: alternative primer sequences are marked “#2”.
