## Supplementary Table S4 for "Stable reference genes for 24-hour circadian profiling of core clock genes in the blood of obstructive sleep apnea patients"

**Supplementary Table S4:** Results of step 1 obtained from RefFinder representing stability values of different algorithms and standard deviation from the BestKeeper algorithm.

| **Rank** | **Delta Ct** | | **BestKeeper** | | **NormFinder** | | **GeNorm** | | **Comprehensive ranking** | |
| --- | --- | --- | --- | --- | --- | --- | --- | --- | --- | --- |
|  | Gene | Average of SD | Gene | SD  (±Cq) | Gene | Stability value | Gene | Stability value | Gene | Geomean o ranking values |
| 1 | *PPIB* | 0.66 | *CDK4* | 0.64 | *ACTB* | 0.097 | *PPIB* | 0.221 | *PPIB* | 2.11 |
| 2 | *PPIB #2* | 0.67 | *PPIA* | 0.75 | *PPIB* | 0.173 | *HPRT1* | 0.221 | *ACTB* | 3.41 |
| 3 | *CDK4 #2* | 0.68 | *RPL13A* | 0.79 | *PPIB #2* | 0.212 | *ACTB* | 0.237 | *PPIB #2* | 3.60 |
| 4 | *TBP* | 0.68 | *CDK4 #2* | 0.86 | *RPL13A* | 0.227 | *PPIB #2* | 0.265 | *CDK4 #2* | 4.53 |
| 5 | *ACTB* | 0.68 | *SDHA* | 0.89 | *SDHA #2* | 0.269 | *CDK4 #2* | 0.306 | *HPRT1* | 4.64 |
| 6 | *RPL13A* | 0.70 | *TBP* | 0.92 | *HPRT1* | 0.274 | *TBP* | 0.332 | *RPL13A* | 4.74 |
| 7 | *HPRT1* | 0.71 | *PPIB #2* | 0.95 | *CDK4 #2* | 0.301 | *RPL13A* | 0.349 | *TBP* | 5.83 |
| 8 | *PPIA* | 0.72 | *SDHA #2* | 0.95 | *TBP* | 0.323 | *PPIA* | 0.362 | *PPIA* | 5.98 |
| 9 | *SDHA #2* | 0.73 | *ACTB* | 1.03 | *GAPDH* | 0.332 | *SDHA* | 0.375 | *CDK4* | 6.45 |
| 10 | *SDHA* | 0.74 | *PPIB* | 1.05 | *PPIA* | 0.383 | *SDHA #2* | 0.383 | *SDHA #2* | 7.75 |
| 11 | *GAPDH* | 0.78 | *HPRT1* | 1.15 | *SDHA* | 0.435 | *GAPDH* | 0.404 | *SDHA* | 8.39 |
| 12 | *CDK4* | 0.85 | *GAPDH* | 1.18 | *CDK4* | 0.568 | *CDK4* | 0.432 | *GAPDH* | 10.69 |
| 13 | *TUBB2A* | 1.76 | *TUBB2A* | 1.31 | *TUBB2A* | 1.598 | *TUBB2A* | 0.616 | *TUBB2A* | 13.00 |
| 14 | *UBC* | 2.98 | *UBC* | 2.53 | *UBC* | 2.929 | *UBC* | 0.953 | *UBC* | 14.00 |

* Abbreviations: SD – standard deviation; Cq – cycle threshold; Alternative primer sequences are marked “*#2*”.
