## Supplementary Table S5 for "Stable reference genes for 24-hour circadian profiling of core clock genes in the blood of obstructive sleep apnea patients"

**Supplementary Table S5:** Results of standard deviations and stability from EndoGeneAnalyzer analysis of the step 2.

| **Gene(s)** | **Individual patient** | | **OSA severity** | | **Time points** | |
| --- | --- | --- | --- | --- | --- | --- |
|  | SD | Stability value | SD | Stability value | SD | Stability value |
| ***PPIB*** | 2.11 | 0.21 | 2.00 | 0.14 | 1.85 | 0.54 |
| ***TBP*** | 1.93 | 0.17 | 1.78 | 0.09 | 1.78 | 0.46 |
| ***ACTB*** | 3.08 | 0.19 | 1.67 | 0.16 | 1.60 | 0.51 |
| ***RPL13A*** | 2.66 | 0.17 | 1.06 | 0.14 | 1.15 | 0.46 |
| ***CDK4 #2*** | 2.10 | 1.39 | 2.12 | 0.95 | 2.09 | 2.04 |
| ***HPRT1*** | 2.20 | 0.31 | 2.09 | 0.24 | 2.03 | 0.69 |

* Abbreviations: SD – standard deviation; Alternative primer sequences are marked “*#2*”.
