## Supplementary Table S6 for "Stable reference genes for 24-hour circadian profiling of core clock genes in the blood of obstructive sleep apnea patients"

**Supplementary Table S6:** Results of step 2 obtained from RefFinder representing stability values of different algorithms and standard deviation from BestKeeper algorithm.

| Rank | **Delta Ct** | | **BestKeeper** | | **NormFinder** | | **GeNorm** | | **Comprehensive ranking** | |
| --- | --- | --- | --- | --- | --- | --- | --- | --- | --- | --- |
|  | Gene | Average of SD | Gene | SD (± Cq) | Gene | Stability value | Gene | Stability value | Gene | Geomean of ranking values |
| 1 | *PPIB* | 3.57 | *RPL13A* | 1.32 | *PPIB* | 1.654 | *HPRT1/TBP* | 0.723 | *TBP/PPIB* | 1.86 |
| 2 | *TBP* | 3.74 | *ACTB* | 1.76 | *TBP* | 2.476 |  |  |  |  |
| 3 | *HPRT1* | 3.80 | *TBP* | 2.20 | *HPRT1* | 2.633 | *PPIB* | 0.785 | *HPRT1* | 2.71 |
| 4 | *CDK4 #2* | 4.87 | *PPIB* | 2.28 | *CDK4 #2* | 3.678 | *CDK4 #2* | 2.012 | *RPL13A* | 3.34 |
| 5 | *RPL13A* | 5.71 | *CDK4 #2* | 2.39 | *RPL13A* | 5.192 | *RPL13A* | 3.974 | *CDK4 #2* | 4.23 |
| 6 | *ACTB* | 5.80 | *HPRT1* | 2.49 | *ACTB* | 5.332 | *ACTB* | 4.582 | *ACTB* | 4.56 |

* Abbreviations: SD – standard deviation; Cq – cycle threshold; Alternative primer sequences are marked “*#2*”.
