## Supplementary Table S7 for "Stable reference genes for 24-hour circadian profiling of core clock genes in the blood of obstructive sleep apnea patients"

**Supplementary Table S7:** Results of standard deviations and stabilities from EndoGeneAnalyzer analysis of the step 3.

| **Gene(s)** | **Individual patient** | | **OSA severity** | | **Time points** | |
| --- | --- | --- | --- | --- | --- | --- |
|  | SD | Stability value | SD | Stability value | SD | Stability value |
| ***PPIB*** | 3.57 | 1.01 | 3.37 | 0.50 | 3.24 | 0.36 |
| ***TBP*** | 3.50 | 1.18 | 2.60 | 0.47 | 2.56 | 0.39 |
| ***ACTB*** | 2.86 | 1.25 | 2.49 | 0.44 | 2.36 | 0.44 |
| ***RPL13A*** | 3.03 | 0.95 | 2.53 | 0.23 | 2.49 | 0.24 |

* Abbreviation: SD – standard deviation.
