## Supplementary Table S8 for "Stable reference genes for 24-hour circadian profiling of core clock genes in the blood of obstructive sleep apnea patients"

**Supplementary Table S8:** Results of step 3 obtained from RefFinder representing stability values of different algorithms and standard deviation from BestKeeper algorithm.

| **Rank** | **Delta Ct** | | **BestKeeper** | | **NormFinder** | | **GeNorm** | | **Comprehensive ranking** | |
| --- | --- | --- | --- | --- | --- | --- | --- | --- | --- | --- |
|  | Gene | Average of SD | Gene | SD (± Cq) | Gene | Stability value | Gene | Stability value | Gene | Geomean of ranking values |
| 1 | *PPIB* | 2.815 | *ACTB* | 1.755 | *PPIB* | 1.709 | *PPIB* | 2.514 | *PPIB* | 1.414 |
| 2 | *ACTB* | 2.886 | *RPL13A* | 2.184 | *ACTB* | 1.855 | *RPL13A* | 2.514 | *ACTB* | 1.661 |
| 3 | *RPL13A* | 2.972 | *TBP* | 2.318 | *RPL13A* | 2.115 | *ACTB* | 2.685 | *RPL13A* | 2.06 |
| 4 | *TBP* | 3.303 | *PPIB* | 2.715 | *TBP* | 2.710 | *TBP* | 2.994 | *TBP* | 3.722 |

* Abbreviations: SD – standard deviation; Cq – cycle threshold.
