## Supplementary Table S9 for "Stable reference genes for 24-hour circadian profiling of core clock genes in the blood of obstructive sleep apnea patients"

**Supplementary Table S9**: The rhythmicity parameters obtained on group-specific cosinor models for each group of participants and for each of the observed genes.

| **Gene** | **Group** | **p** | **q** | **Amplitude** | | | **Mesor** | | | **Acrophase** | | | **Acrophase**  **[h]** |
| --- | --- | --- | --- | --- | --- | --- | --- | --- | --- | --- | --- | --- | --- |
|  |  |  |  | **Value** | **p** | **q** | **Value** | **p** | **q** | **Value** | **p** | **q** |  |
| ***BMAL1*** | Control | 0.3607 | 0.5900 | 0.071 | 0.2988 | 0.4555 | 0.060 | 0.1162 | 0.1550 | -3.642 | n.s. | n.s. | 13.911 |
|  | Mild | 0.4188 | 0.5900 | 0.024 | 0.1920 | 0.4555 | 0.026 | 0.0189 | 0.1138 | -5.456 | n.s. | n.s. | 20.841 |
|  | Moderate | 0.3207 | 0.5900 | 3.076 | 0.3513 | 0.4555 | 1.433 | 0.3304 | 0.3304 | -2.952 | n.s. | n.s. | 11.276 |
|  | Severe | 0.8942 | 0.8942 | 0.024 | 0.6613 | 0.6613 | 0.089 | 0.0983 | 0.1474 | -5.376 | n.s. | n.s. | 20.536 |
| ***CRY1*** | Control | 0.1780 | 0.5900 | 0.085 | 0.2136 | 0.4555 | 0.102 | 0.0391 | 0.1474 | -3.648 | n.s. | n.s. | 13.934 |
|  | Mild | 0.3214 | 0.5900 | 0.616 | 0.3785 | 0.4555 | 0.358 | 0.2357 | 0.2571 | -5.208 | n.s. | n.s. | 19.892 |
|  | Moderate | 0.6586 | 0.7873 | 11.203 | 0.3461 | 0.4555 | 0.160 | 0.0694 | 0.1474 | -3.404 | n.s. | n.s. | 13.001 |
|  | Severe | 0.3698 | 0.5900 | 0.049 | 0.3796 | 0.4555 | 0.276 | 0.0904 | 0.1474 | -4.259 | n.s. | n.s. | 16.269 |
| ***PER2*** | Control | 0.4425 | 0.5900 | 0.060 | 0.1909 | 0.4555 | 0.079 | 0.0554 | 0.1474 | -3.903 | n.s. | n.s. | 14.911 |
|  | Mild | 0.3653 | 0.5900 | 0.438 | 0.2710 | 0.4555 | 0.324 | 0.0968 | 0.1474 | -5.422 | n.s. | n.s. | 20.710 |
|  | Moderate | 0.7217 | 0.7873 | 1.023 | 0.4799 | 0.5236 | 1.480 | 0.1590 | 0.1908 | -3.358 | n.s. | n.s. | 12.828 |
|  | Severe | 0.4407 | 0.5900 | 0.036 | 0.2107 | 0.4555 | 0.137 | 0.0067 | 0.0809 | -5.809 | n.s. | n.s. | 22.192 |

* Abbreviation: n.s. – non-significant.
