## Supplementary Table S10 for "Stable reference genes for 24-hour circadian profiling of core clock genes in the blood of obstructive sleep apnea patients"

**Supplementary Table S10**: The statistically significant rhythmicity parameters obtained on personalized cosinor models for each individual participant and for each of the observed genes.

| **Gene** | **Patient ID** | **Group** | **Amplitude** | **p (amplitude)** | **Acrophase** | **Acrophase [h]** |
| --- | --- | --- | --- | --- | --- | --- |
| ***BMAL1*** | P1 | Control | 0.00425 | 4.66553E-11 | -1.11632673 | 4.26405403 |
|  | P2 | Control | 0.060191 | 7.46845E-06 | 2.434581991 | 14.7005818 |
|  | P3 | Mild | 0.085935 | 0 | 0.541208726 | 21.93273495 |
|  | P4 | Mild | 0.029186 | 0.00010875 | -0.94427217 | 3.606854013 |
|  | P5 | Mild | 0.009471 | 0.010991195 | 2.404003869 | 14.81738162 |
|  | P6 | Mild | 0.123492 | 0.038149372 | 1.105492929 | 19.77732806 |
|  | P7 | Moderate | 0.005344 | 0.000310015 | 2.977527063 | 12.62668439 |
| ***CRY1*** | P8 | Control | 0.011067 | 3.93393E-09 | -0.15638869 | 0.597360799 |
|  | P9 | Control | 0.008767 | 0.008898627 | 1.9408773 | 16.58639481 |
|  | P2 | Control | 0.262639 | 0.012280493 | 2.783128226 | 13.36923325 |
|  | P10 | Control | 0.024815 | 0.015744202 | 0.727688855 | 21.22043332 |
|  | P11 | Mild | 0.102313 | 4.6574E-125 | -0.12461348 | 0.475988444 |
|  | P4 | Mild | 0.040103 | 2.28518E-07 | -1.11083241 | 4.243067265 |
|  | P5 | Mild | 0.025212 | 5.04847E-07 | 1.743656182 | 17.33972399 |
|  | P6 | Mild | 0.086707 | 0.037089256 | 1.301938626 | 19.02696077 |
|  | P12 | Severe | 0.016704 | 7.58688E-63 | 2.242394766 | 15.43468293 |
|  | P13 | Severe | 0.018096 | 1.66146E-05 | 0.035646443 | 23.86384062 |
|  | P14 | Severe | 0.066099 | 0.012479956 | 2.55428976 | 14.24333181 |
|  | P15 | Severe | 0.050427 | 0.045094712 | -0.4871363 | 1.860723589 |
|  | P16 | Severe | 0.272777 | 0.047325607 | 1.175803791 | 19.50876035 |
| ***PER2*** | P4 | Mild | 0.009401 | 2.9747E-08 | -1.25363778 | 4.788543578 |
|  | P17 | Mild | 1.215952 | 0.018982656 | -0.38822155 | 1.482897095 |
|  | P18 | Mild | 0.003813 | 0.047919036 | -1.73827654 | 6.639727278 |
|  | P7 | Moderate | 0.00475 | 6.06836E-06 | 0.793181601 | 20.97026946 |
|  | P19 | Severe | 0.229097 | 0.002272685 | -0.02237897 | 0.085481365 |
