## Supplementary Table S11 for "Stable reference genes for 24-hour circadian profiling of core clock genes in the blood of obstructive sleep apnea patients"

**Supplementary Table S11:** Number of patients with statistically significant rhythmicity of core clock genes across OSA severity groups.

| **Gene** | **OSA severity** | **Number of patients** | **SUM** |
| --- | --- | --- | --- |
| ***BMAL1*** | Control | 2 | 7 |
|  | Mild | 4 |  |
|  | Moderate | 1 |  |
| ***CRY1*** | Control | 4 | 13 |
|  | Mild | 4 |  |
|  | Severe | 5 |  |
| ***PER2*** | Mild | 3 | 5 |
|  | Moderate | 1 |  |
|  | Severe | 1 |  |
